# Mining genes underlying perennial-to-annual transitions in *Oryza* with comparative genomics

**DOI:** 10.1101/2025.09.01.673603

**Authors:** Zhenhua Lu, Yan Zhou, Guangzhao Yang, Laura Ellen Rose, Fengyi Hu, Zheng Li

## Abstract

Perennial rice has shown great potential for alleviating multidimensional sustainability issues that are prevalent in current rice agriculture, prompting widespread interest in its breeding. Currently, one major hurdle for these breeding efforts is our scarce knowledge about the evolutionary mechanisms of the perennial-to-annual transition in *Oryza*. Here, by performing comparative genomics of three annual-perennial pairs of *Oryza* species (*Oryza rufipogon*/*O. nivara*, *O. glumaepatula*/*O. glaberrima*, and *O. meyeriana*/*O. brachyantha*), we identified 91 gene families that are present in all the perennial species but are missing in any of the annual species. Annotations and functional prediction of these gene families demonstrated that they may be associated with a battery of physiological components of perenniality, such as phenology, energy allocation, and source–sink dynamics. Furthermore, comparative co-expression analysis revealed that many of these families may also play important regulatory roles for orchestrating diverse processes that differ between perennial and annual *Oryza* species, including reproduction, root development, and response to environmental stimuli. These results highlight the potential complexity of the emergence of annuality in the *Oryza* genus and offer promising candidates for future functional studies.

## Background

Our current cultivated rice species are annual plants that are theorised to be directly or indirectly domesticated from their wild perennial progenitors (Kovach et al. 2007; Sang and Ge 2007). While this perennial-to-annual transition is usually not considered to be part of the domestication syndrome, it intertwines with many agronomic changes favoured by our ancestors (e.g., larger grains and higher yield) due to the distinct resource allocation strategies of annuals and perennials (Chen et al. 2019). Unlike perennial plants, whose assimilated photosynthetic energy is divided between sexual reproduction and perennating structure development, an annual plant channels nearly all its resources into seeds over the course of a growing season (Li et al. 2022). The high-yielding nature of annual cultivated rice has given rise to the one-sow, one-harvest routine of modern rice agriculture and underpins the livelihood of half of the world’s population who eat rice as the primary food source. However, multiple sustainability issues associated with this agricultural practice have been gradually recognised. In particular, annual cropping causes long-standing soil disruption and demands significant energy and labour input (Cox et al. 2006). Facing these issues, considerable efforts have been put in place to breed perennial rice, and its recent landmark breakthrough may usher in a new landscape of future sustainable agriculture (Zhang et al. 2023). Despite this promising prospect, further development of perennial rice is hindered by our inadequate evolutionary and molecular understanding of the perennial-to-annual transition in *Oryza*.

The *Oryza* genus, consisting of 24 species of wide genetic diversity, is regarded as an ideal model for studying plant evolution (Vaughan et al. 2003). However, the evolutionary pattern from perennial to annual species is much less understood in comparison with other domestication-related traits (Vaughan et al. 2008). Apart from the two cultivated species, only a few wild *Oryza* species have an annual life history, indicating that the perennial habit is the ancestral life history for the genus (Kellogg 2009). Phylogenetic evidence suggests that the annual habit has evolved independently among AA genome species (*O. barthii*, *O. glaberrima*, *O. meridionalis*, and *O. nivara*), BB genome species (*O. punctata*), and FF genome species (*O. brachyantha*) (Menguer et al. 2017). These annual species are distributed across continents, and they often share habitats with perennial ones (Vaughan et al. 2003; Sengupta and Majumder 2010). Strikingly, both annual and perennial individuals of *O. rufipogon*, which is generally considered a perennial species, have been observed in the same populations (Sano et al. 1980). Hence, it seems that there are no universal environmental factors driving this perennial-to-annual evolution, implying that multiple and complex genetic mechanisms are involved.

Gene loss, which broadly refers to changes that eliminate the function of a gene, has been recognised as a universal and frequent evolutionary event that can prompt adaptive phenotypic diversity across all life kingdoms (Albalat and Cañestro 2016). In plants, one well-documented example of the adaptive loss-of-function has been reported in changes in flower petal colour that led to new pollination syndromes in *Ipomoea* (Zufall and Rausher 2004). Some species of this genus experienced an adaptive change from blue to red flowers due to the loss of the flavonoid 3’-hydroxylase gene, inducing the shift from bee pollination to bird pollination. In *Oryza*, the potential importance of gene loss in the perennial-to-annual transition is hinted by the fact that the hybrid offspring between the annual *O. sativa* and the perennial *O. longistaminata* tend to have perennial genotypes (Hu et al. 2003). Moreover, extensive gene loss has been identified between annual species and perennial species in the *Arabidopsis* genus (Hu et al. 2011), and more specifically, loss-of-function mutations of floral repressor genes are directly associated with the diversification of perennial and annual flowering behaviour in the Brassicaceae family (Li et al. 2024; Zhai et al. 2024).

Given these clues, in this study we focused on gene loss events that potentially underpin the life history switch from perennial to annual in *Oryza*. Comparative genomics analysis of three annual-perennial pairs revealed gene families that may contribute to the perennial life history but are lost in annuals. Further functional and evolutionary insights were obtained from synteny, expression, motif, and comparative co-expression network analysis. These results provide candidates for future functional studies and may support molecular breeding efforts of perennial rice.

## Results and Discussion

### Identification of candidate genes

Three pairs of perennial/annual sister species were used in our comparative genomics analysis, namely, *O. rufipogon* (perennial) and *O. nivara* (annual), *O. glumaepatula* (perennial) and *O. glaberrima* (annual), and *O. meyeriana* (perennial) and *O. brachyantha* (annual) (Table 1). These species were selected based on the quality and availability of their genome sequences and their phylogenetic positions. All these genome assemblies were of high quality, with an N50 length ranging from 11.4 to 64.5 Mb and a coverage of 102 × to 168 ×. The phylogenetic positions of these species were previously reported (Kellogg 2009; Zhang et al. 2014; Stein et al. 2018). We selected these pairs following the criteria that the species in each pair are as closely related to each other as possible; thus, their genotypic differences can to the largest degree be attributed to genes responsible for the changes in life history. A phylogenetic tree was constructed to specifically illustrate the evolutionary relationship between the selected species (Figure 1A). The tree was inferred from 15,559 multi-copy or single gene families, and it is in agreement with other published phylogenies for the *Oryza* genus (Kellogg, 2009; Zhang et al. 2014; Stein et al. 2018).

**Table 1.**
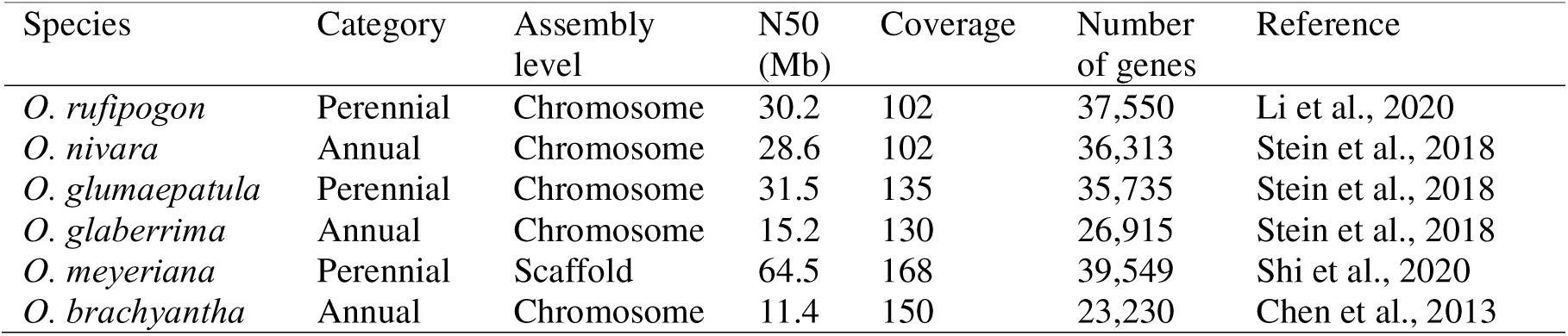
Genome information of the six *Oryza* species in this study.

| Species | Category | Assembly level | N50 (Mb) | Coverage | Number of genes | Reference |
| --- | --- | --- | --- | --- | --- | --- |
| *O. rufipogon* | Perennial | Chromosome | 30.2 | 102 | 37,550 | Li et al., 2020 |
| *O. nivara* | Annual | Chromosome | 28.6 | 102 | 36,313 | Stein et al., 2018 |
| *O. glumaepatula* | Perennial | Chromosome | 31.5 | 135 | 35,735 | Stein et al., 2018 |
| *O. glaberrima* | Annual | Chromosome | 15.2 | 130 | 26,915 | Stein et al., 2018 |
| *O. meyeriana* | Perennial | Scaffold | 64.5 | 168 | 39,549 | Shi et al., 2020 |
| *O. brachyantha* | Annual | Chromosome | 11.4 | 150 | 23,230 | Chen et al., 2013 |

**Figure 1.**
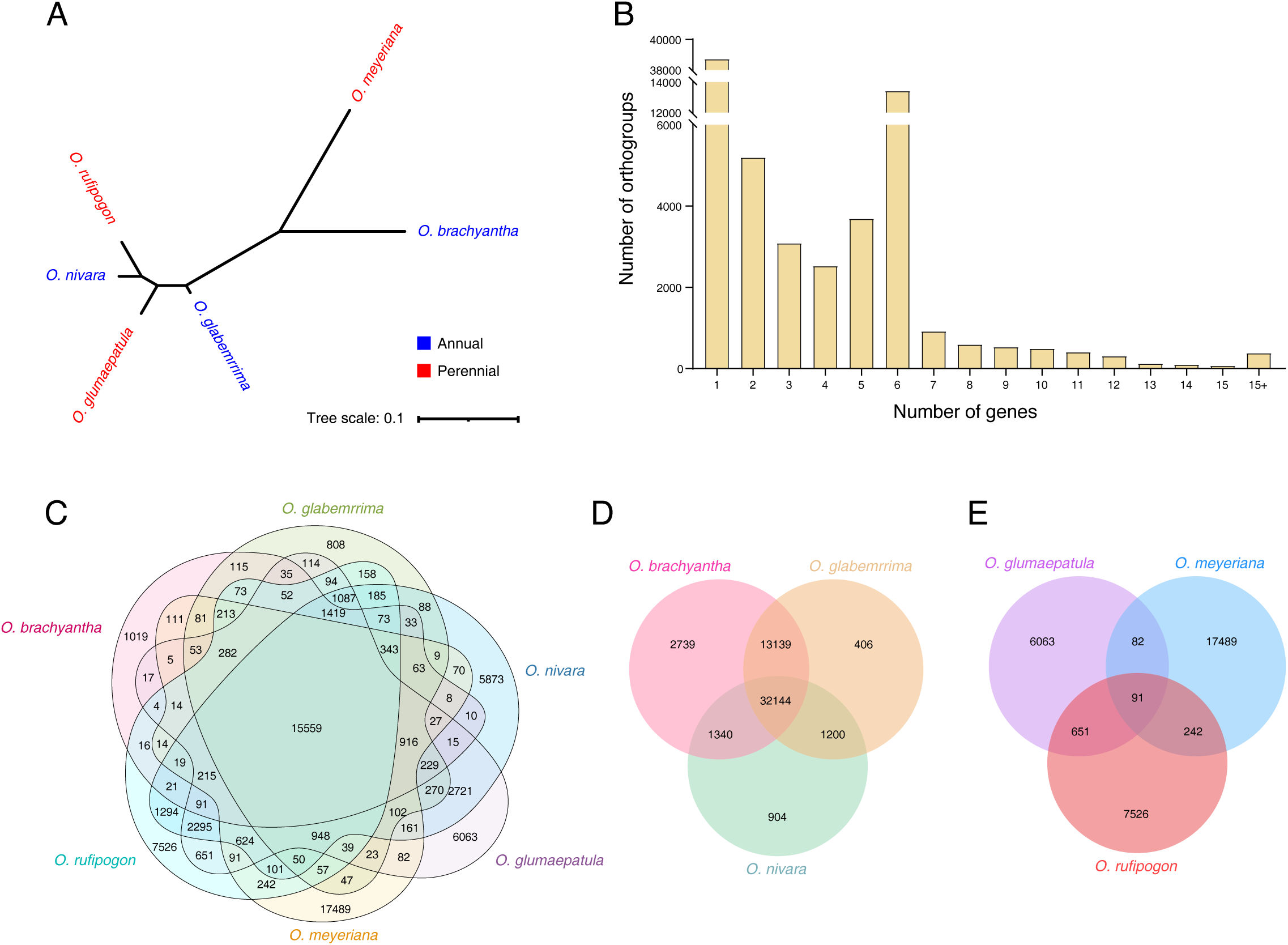
Identification of candidate genes underlying perennial-to-annual transitions in *Oryza*. **A** Phylogenetic tree of the six *Oryza* species. The tree was inferred from 15,559 multi-copy or single gene families using the built-in STAG algorithm in OrthoFinder, and the scale bar indicates substitutions per site. **B** OrthoFinder result of sorting genes of the six *Oryza* species into gene families. **C** Venn diagram of all gene families. **D** Venn diagram of gene families that were lost in at least one annual *Oryza* species. **E** Venn diagram of gene families that are present in at least one perennial *Oryza* species but were lost in all annual *Oryza* species.

To find genes that have been lost or undergone pseudogenization in annual *Oryza* species, the protein sequences from the selected species were first sorted into gene families using OrthoFinder (Emms and Kelly 2019). We obtained a total of 70,507 gene families, the majority of which contained fewer than seven genes (Figure 1B; Supplementary Table 1). Except for gene families containing one single orphan gene, gene families with six genes accounted for the highest proportion, with a total of 13,423 families. Figure 1C visualizes the distribution of gene families among the species. A total of 15,559 gene families are shared by the selected species in this study, likely representing core rice genes (Shang et al. 2022). Of all detected gene families, 32,144 families contain at least one gene from a perennial species but missing genes from all annual species (Figure 1D). These gene families were further investigated, and their distribution is illustrated in Figure 1E. A large number of these non-annual-gene families were found to be present in only one perennial species—6063 in *O. glumaepatula*, 17,489 in *O. meyeriana*, and 7526 in *O. rufipogon*. By contrast, when two perennial species were considered, the quantity of genes was significantly shrunk, with a range between 173 to 742. Finally, the intersection of all three perennial species contains 91 families. These ones are candidate families with potential importance for the transition from perennial to annual as they encompass at least one gene from each perennial species and have no genes from annuals.

Compared with a similar comparative genomics study conducted for Brassicaceae species (Heidel et al. 2016), a much larger number of differential gene families between perennial and annual species were identified in this study. This is not surprising since the species pairs used in the Brassicaceae study were very closely related and show little morphological difference other than life history. By contrast, as annual species are sparse in the *Oryza* genus, the annual/perennial pairs studied in this study are more distant and exhibit many interspecific differences in addition to life history (Stein et al. 2018). Even for *O. rufipogon* and *O. nivara*, two relatives that are sometimes regarded as ecotypes of one species, over a dozen of phenotypic traits (e.g., anther length, culm length, and panicle shape) have been identified as species-distinguishing traits (Cai et al. 2019; Zheng and Ge 2010), which are underpinned by significantly differing genetic architectures (Li et al. 2020; Meng et al. 2024). Nevertheless, when more than one perennial species was considered, the number of differential families were greatly reduced (Figure 1E), indicating that there is a core set of genes underlying the evolution of annuality in *Oryza*.

### Functional overview of the candidate genes

The 91 candidate families include a total of 400 genes from the three perennial species. To gain more functional insights from these genes, we performed EggNOG annotation using their protein sequences (Supplementary Table 2). In line with a previous study finding gene loss events between perennials and annuals in the Brassicaceae (Heidel et al. 2016), many of our candidate genes were predicted to encode signal transduction-related proteins (e.g., F-box proteins, Myb/SANT-like DNA-binding proteins, and kinases). To date, the most well-understood genetic basis for the evolution of annuals from perennials concerns their differential flowering behaviours (Wang et al. 2009; Li et al. 2024; Zhai et al. 2024). Several floral repressor genes have been found to dictate perennial or annual flowering behaviour in a dosage-dependent manner, and one main determinant here is their distinct stabilities of vernalisation-induced gene repression between perennials and annuals. Previous studies have implied that this dissimilarity may stem from differences in upstream signal transduction pathways (Wang et al. 2009; Coustham et al. 2012; Kemi et al. 2013; Hepworth et al. 2020; Zhai et al. 2024). Although rice plants do not require vernalisation for floral induction, components of the vernalisation pathway known for *Arabidopsis* are involved in photoperiod regulation of flowering in rice (Wang et al. 2013). Hence, some of these signal transduction-related genes in perennial *Oryza* species may be associated with photoperiod-mediated perennial flowering behaviour. In addition, a number of genes from the candidate families were annotated to be disease resistance protein-encoding genes. While change in disease resistance has not been considered as an element of the perennial-to-annual transition, such a reduction of disease-resistance gene repertoire perhaps represents a secondary evolutionary outcome of short-lived annual life history on the genome—annual plants are exposed to fewer pathogens during their short life cycle, and thus they further modulate resources for disease resistance capacity to reproduction (Friedman 2020).

Further functional insights can be gained from the Gene Ontology (GO) and Kyoto Encyclopedia of Genes and Genomes (KEGG) terms associated with the candidate gene families (Figures 2A and B; Supplementary Table 3). In addition to many signal transduction-related terms (e.g., ‘cell periphery’, ‘protein modification process’, ‘hormone metabolic process’, ‘plant hormone signal transduction’, ‘PPAR signalling pathway’, and ‘cAMP signalling pathway’), a number of terms are connected with energy metabolism, such as ‘mitochondrion’, ‘ATPase-coupled transmembrane transporter activity’, ‘ATP hydrolysis activity’, ‘fatty acid metabolism’, and ‘peroxisome’. As mentioned earlier, annuals and perennials differ significantly in their energy allocation strategies. A comparative study of perennial and annual *Lupinus* species revealed that, in comparison with the perennial *Lupinus variicolor*, the annual *L. nanus* triples its energy investment ratio between reproduction and vegetative growth (Pitelka 1977). Although such a resource use trade-off has been a central topic of plant physiology (Doust 1989), little is known about the genetic basis of this strategic difference between annuals and perennials. Our results suggest that the loss of energy metabolism-related genes may contribute to the reproduction-centred energy allocation strategy in annual *Oryza* species, yet the detailed mechanisms of how these genes are involved are worth further investigation.

**Figure 2.**
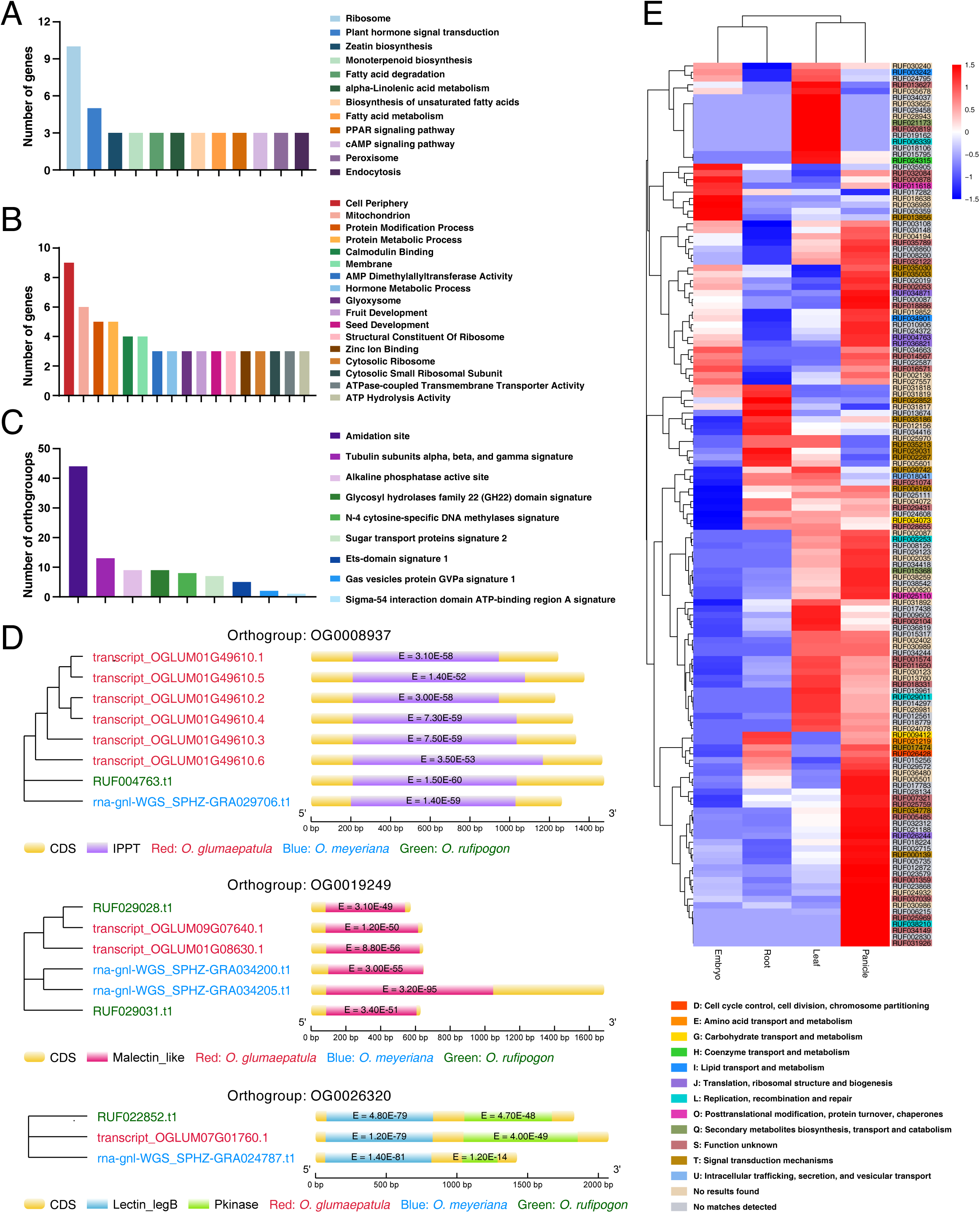
Functional overview of the candidate genes. **A** GO terms associated with the candidate genes. **B** KEGG pathways associated with the candidate genes. **C** Meme motif analysis result of the candidate genes. **D** Top high-confidence Pfam domains detected in the candidate genes. **E** Expression level of the candidate genes in *O. rufipogon*. Colour scale represents Zscore of scaled CPM normalized expression values. Gene IDs are coloured according to Clusters of Orthologous Gene (COG) categories generated by EggNOG annotation.

To look into the precise action of mode of these candidate gene families, we also performed motif enrichment and Pfam analyses. One predominant motif uncovered by the MEME suite is amidation site (Figure 2C; Supplementary Table 4). The amidation process is known to be involved in multiple aspects of hormone-related signalling, such as hormone homeostasis and biosynthesis (Mano et al. 2010; Li et al. 2018; Široká et al. 2024). In particular, strigolactone activation by amidation has been considered to be a link between branching control and nitrogen stress response in *Arabidopsis* (Zhu and Kranz, 2012). Interestingly, both branching and nitrogen use efficiency are standing traits that experienced dramatic changes during the evolution of *Oryza* species (Chen et al. 2019; Hu et al. 2023), thereby implying the potential importance of amidation also in life history transitions. Consistent with the previous discussion on the potential signalling roles of the candidate families, high-confidence Pfam domains include ‘malectin_like’, ‘lectin_legB’, ‘isopentenyl pyrophosphate transferase (IPPT)’, and ‘pkinase’ (Figure 2D; Supplementary Table 5), which are all implicated in the signalling pathways of various hormones (Lindner et al. 2014; Bellande et al. 2017; Feng et al. 2022; Park and Yoon 2025).

To corroborate these functional insights, the expression levels of all the identified candidate genes from *O. rufipogon* are plotted in Figure 2E. In agreement with the differential flowering and reproductive strategies between annual and perennial *Oryza* species, the majority of the candidate genes were found to be highly expressed in panicles. Interestingly, several genes annotated with the term ‘signal transduction mechanisms’ were detected to exhibit high expression levels in roots, and, to a lesser degree in leaves. One distinguishing physiological feature of the perennial versus the annual is the former’s source– sink dynamics between leaves and roots, both of which can serve as source and sink organs. During the life cycle of some perennials, nutrients can be relocated from photosynthetic tissues to the roots for storage under non-optimal growth conditions and then recycled to the leaves for subsequent growth when favourable conditions return (Yang et al. 2016). Our knowledge regarding the regulatory mechanism of this dynamic pattern is scarce. One hypothesised scenario implied by our results is that these signal transduction-related genes may orchestrate such dynamic patterns, and that the loss of these genes may lead to rewiring of the network, ultimately resulting in the depletion of this recycling mechanism in annuals.

### Comparative co-expression network analysis

It is well established that functionally related genes tend to be transcriptionally coordinated (i.e., co-expressed), and thus co-expression networks are a powerful tool to reveal relationships between genes and generate hypotheses on gene function (Mutwil et al. 2021). Recently, comparison of gene co-expression networks among species has emerged as an approach valuable for the study of evolution (Ruprecht et al. 2017; Ovens et al. 2021). Here, to unravel the potential regulatory roles and the evolutionary significance of the candidate families in developmental transitions associated with alternative life histories, we performed a comparative analysis of gene co-expression networks. Given the wide availability of functional genomics knowledge in annual cultivated rice and limited gene expression data for most wild rice species, a gene co-expression network was first constructed for the perennial species *O. rufipogon* (Supplementary Table 6), and it is compared with a previously published gene co-expression network constructed for the annual species, *O. sativa* (Zhang et al. 2022).

A total of 48 modules were obtained from the *O. rufipogon* co-expression network (Supplementary Table 6). Two modules that contain the largest number of candidate genes (both >20), termed Module 1 and Module 2, were selected to investigate further; all other modules each contain no more than four candidate genes and were thus deemed to be insignificant. For both of these *O. rufipogon* modules, genes homologous to *O. sativa* genes were mapped (referred to as homologous genes hereafter), and their belonging modules in the *O. sativa* co-expression network were determined. In Module 1, 22 candidate genes identified through our comparative genomics analysis are highly interconnected with homologous genes from seven major *O. sativa* modules (modules containing only a few homologous genes are not discussed here) (Figure 3; Supplementary Table 7). Of note, these homologous genes are largely located in different modules within the *O. sativa* co-expression network, and therefore the corresponding networks of *O. sativa* are more fragmented than for *O. rufipogon*. Gene loss along annual lineages may underlie the disappearance of connections between these modules and may partially reflect the evolutionary mechanism of annuality in *Oryza* species. TO and GO annotations of these seven *O. sativa* modules (i.e., M0010, M0086, M0016, M0027, M0050, M0030, and M0034) were used to probe into the potential regulatory roles of the candidate genes. Apart from several modules associated with the above-mentioned processes, such as the reproductive process (also including ‘seed development trait’ and ‘reproductive shoot system development’) and energy metabolism (including ‘starch content’ and ‘lipid metabolic process’), two modules of particular interest are M0034 and M0086, whose genes are enriched in the terms of ‘root development’ and ‘plant-type cell wall organization’, respectively. While the anatomical differences in roots have not been comprehensively compared between annual and perennial *Oryza* species, research on other plant species demonstrated that perennial plants generally have a root system conductive to nutrient uptake and longevity, typically with higher tissue density, diameter, and dry weight (Craine et al. 2001; Roumet et al. 2006; Monti and Zatta 2009). The *O. sativa* module M0086 is characterised by several expansin genes, which are essential for both vegetative and reproductive growth (Choi et al. 2003). Taken together, Module 1 highlights the possible regulatory roles of our identified candidate genes on coordinating resource acquisition and allocation, as well as their resultant growth and developmental strategies.

**Figure 3.**
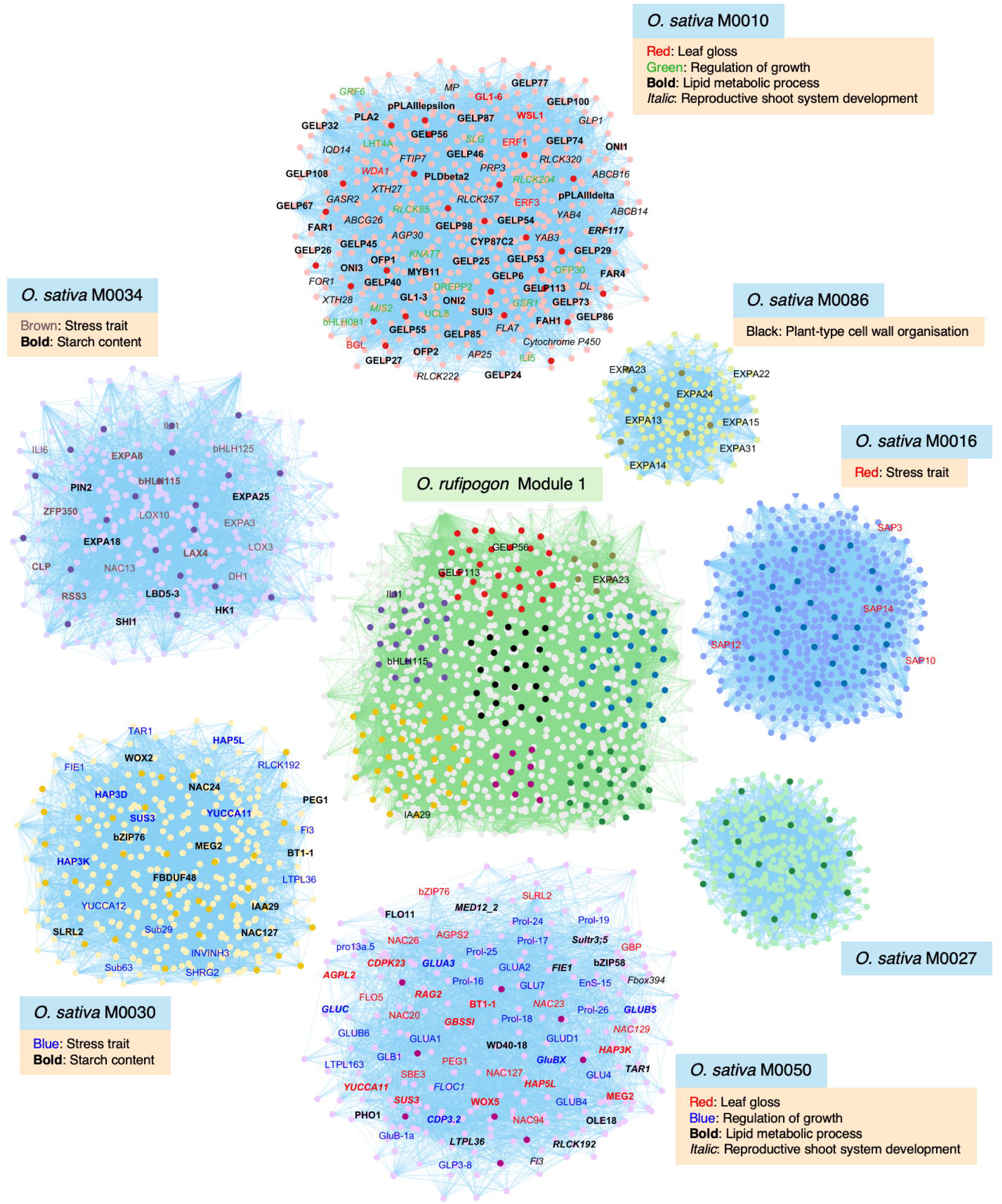
Comparative co-expression analysis between *O. rufipogon* Module 1 and the co-expression network for *O. sativa*. In *O. rufipogon* Module 1, the identified candidate genes are represented by black nodes in the middle and genes homologous to *O. sativa* genes in major *O. sativa* co-expression modules highlighted by colour. in *O. sativa* modules (M0010, M0086, M0016, M0027, M0050, M0030, and M0034), genes homologous to the *O. rufipogon* genes in *O. rufipogon* Module 1 are highlighted in the same colour as their corresponding *O. rufipogon* genes, and gene names are shown for genes with selected significant enriched GO and TO terms that are potentially associated with perennial-to-annual transitions.

In *O. rufipogon* Module 2, 20 candidate genes are linked with homologous genes from three major *O. sativa* modules, namely, M0016, M0030, and M0013 (Figure 4; Supplementary Table 8). Similar to Module 1, these candidate genes in Module 2 may also regulate processes pertaining to seed development (homologous genes from M0030) and stress responses (homologous genes from M0016). On top of these two gene groups, a large number of homologous genes from M0013 are involved in response to environmental stimuli. Overall, Module 2 perhaps depicts the complex interactions between plants and the environment. It is assumed that, when nascent annuals evolved from perennial ancestors, their tolerance to abiotic stressors would attenuate as they faced fewer seasonal extremes than their perennial ancestors (Lundgren and Des Marais 2020). Hence, the loss of the candidate genes in Module 2 may represent genomic changes associated with different adaptive responses following life history transitions.

**Figure 4.**
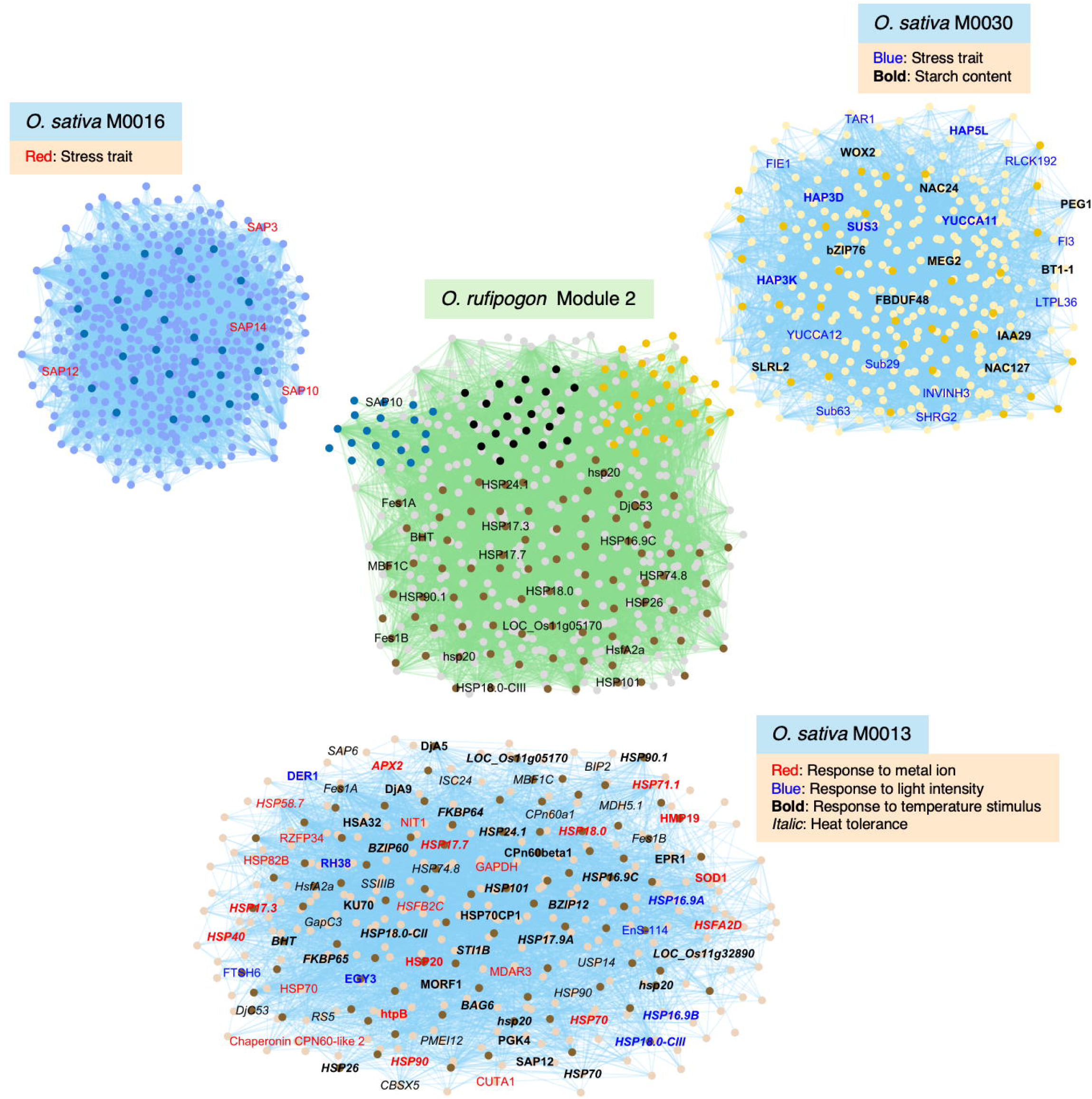
Comparative co-expression analysis between *O. rufipogon* Module 2 and the co-expression network for *O. sativa*. In *O. rufipogon* Module 2, the identified candidate genes are represented by black nodes in the middle and genes homologous to *O. sativa* genes in major *O. sativa* co-expression modules highlighted by colour. in *O. sativa* modules (M0030, M0013, and M0016), genes homologous to the *O. rufipogon* genes in *O. rufipogon* Module 1 are highlighted in the same colour as their corresponding *O. rufipogon* genes, and gene names are shown for genes with selected significant enriched GO and TO terms that are potentially associated with perennial-to-annual transitions.

### Evolutionary analysis

We next performed synteny analysis to shed light on the evolutionary pattern of the missing genes in annual *Oryza* species. The perennial species *O. rufipogon* and its annual descendant *O. nivara* were included in this analysis because these are the most closely related annual-perennial species in our study. This ensures that additional evolutionary changes do not further confound our ability to ascertain candidates involved in the evolutionary shift between perenniality and annuality. Of the 152 candidate genes from *O. rufipogon*, 48 pseudogenes were predicted in *O. nivara*, accounting for around 31.58% of the total. The genomic locations of these pseudogenes are displayed in Figure 5A (Supplementary Table 9). The cases where no pseudogene could be identified likely represent the presence/absence variation of the genes between *O. rufipogon* and *O. nivara*.

**Figure 5.**
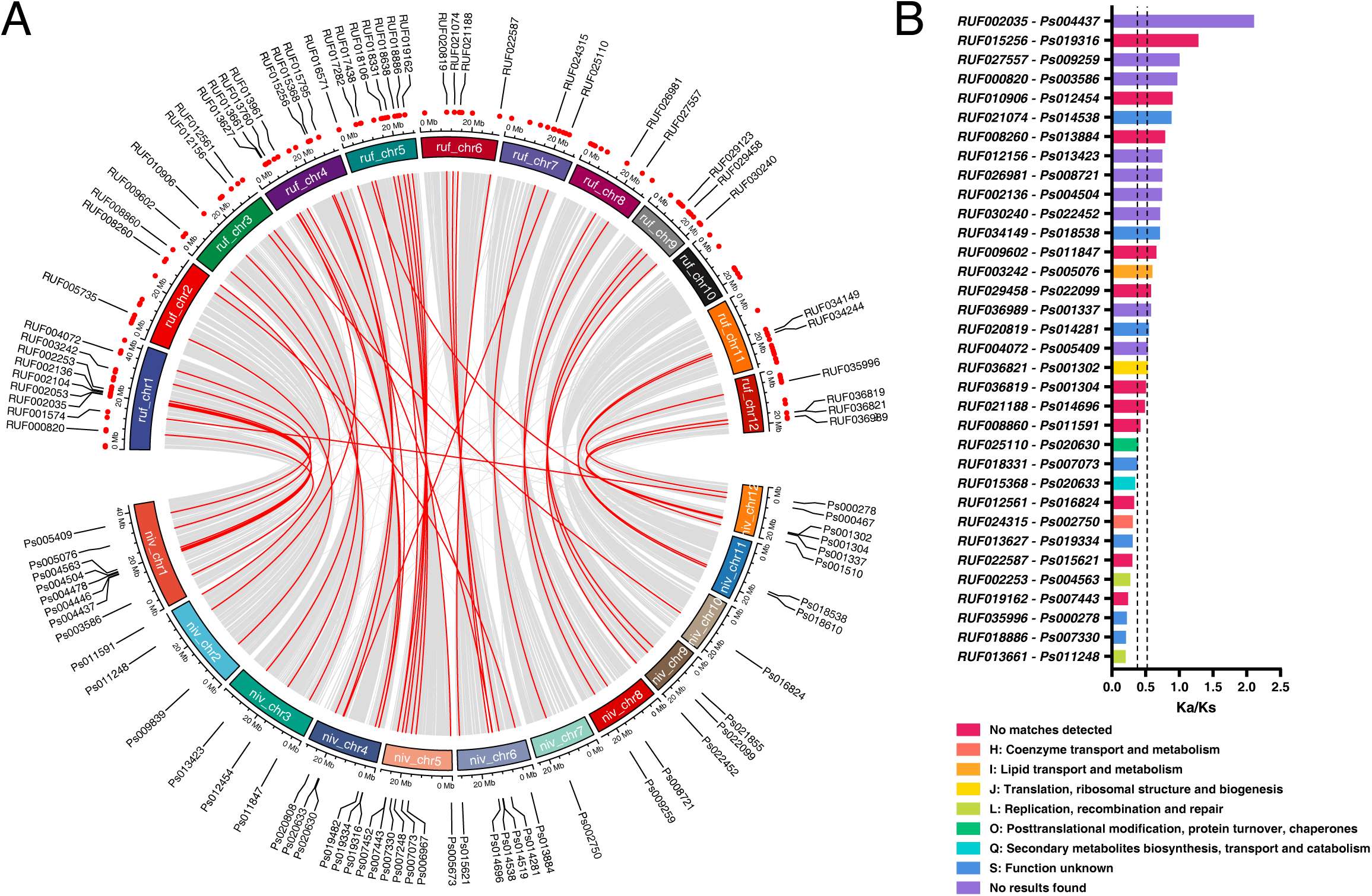
Evolutionary analysis of the candidate genes from *O. rufipogon*. **A** Circos diagram of *O. rufipogon* and *O. nivara* chromosomes showing the candidate genes and their corresponding pseudogenes. Gene IDs are only shown for *O. rufipogon* candidate genes (represented by red dots) for which *O. nivara* pseudogenes (labelled with a pseudogene ID generated by PseudoPipe) were detected. **B** Ka/Ks values of gene–pseudogene pairs. Dashed lines represent the upper and lower bounds of the 95% confidence interval calculated from a set of 16,362 families in which at a minimum contain single genes from *O. rufipogon* and *O. nivara*. Bars are coloured according to COG categories generated by EggNOG annotation.

To estimate the date of pseudogenisation, the Ka/Ks ratios were calculated for the *O. rufipogon* candidate genes and their corresponding pseudogenes in *O. nivara.* These ratios were compared with the Ka/Ks ratio range of a set of 16,362 families in which at a minimum contain single genes from *O. rufipogon* and *O. nivara*. We found 19 pseudogenes whose Ka/Ks ratios were larger than the upper bound of the 95% confidence interval, whereas 10 pseudogenes showed a ratio lower than the lower bound (Figure 5B). Genes belonging to the group in the upper extreme of the Ka/Ks values may indicate old pseudogenisation events, while those genes in the lower extreme of the Ka/Ks values may be more recent events. Furthermore, purifying selection presumably still acts on the perennial gene after the pseudogenisation; thus, recent pseudogenisation in the annual species may result in a low Ka/Ks ratio. According to our analysis, the 19 pseudogenes with a high Ka/Ks ratio may represent key evolutionary events for early diversification of perennial/annual transition caused by the changing environment. Following these changes, the 10 low Ka/Ks pseudogenes were likely formed more recently largely due to purifying selection. As shown in Figure 5B, early pseudogenisation may relate to the processes of ‘lipid transport and metabolism’ and ‘translation, ribosomal structure and biogenesis’, while recent pseudogenisation may have impact on ‘secondary metabolites biosynthesis, transport and catabolism’, ‘coenzyme transport and metabolism’, and ‘replication, recombination and repair’. Nonetheless, as most our detected gene/pseudogene pairs are not fully functionally annotated, investigation of the detailed biological significance of these genes might be of future research interest.

## Conclusions

Since we are now confounded with an expanding array of sustainability issues in rice cultivation, the multiple benefits of perennial rice have sparked considerable interests in its breeding research. There are two general future directions: one is to enhance the perenniality of existing perennial rice lines, and the other is to take advantage of genome editing technology to perennilise elite rice varieties or *de novo* efforts to domesticate perennial wild rice species. Importantly, both of these directions necessitate an in-depth genetic and molecular understanding of perenniality. In this study, we identified gene families that could potentially have impact on annual/perennial transitions in *Oryza*. In contrast to previous reports supporting the notion that the genetic differences between perennials and annuals are rather minor (Melzer et al. 2008; Heidel et al. 2016; Zhai et al. 2024), the relatively large number of candidate gene families suggests that these transitions may entangle intricate genetic mechanisms. Indeed, our comparative co-expression analysis revealed that the loss of these genes might make alterations to diverse biological processes underlying perenniality. Our results therefore bear out the assertion that perenniality and annuality are complex syndromes (Lundgren and Des Marais 2020) and imply that a wide range of aspects of perenniality need to be considered in a combinatorial manner for improving the overall agronomic performance of perennial rice.

## Materials and Methods

### Genome assembly and annotation data

The draft genome assembly and genome annotation of *O. rufipogon* were downloaded from the National Genomics Data Centre (https://ngdc.cncb.ac.cn/) under the accession number PRJCA002346. The genome assembly and genome annotation of *O. brachyantha* was obtained from http://www.gramene.org/Oryza_brachyantha. For *O. meyeriana*, its genome assembly and annotation were downloaded from BIG Genome Warehouse (accession number: GWHAAKB00000000) and the Figshare database (DOI: 10.6084/m9.figshare.8191316), respectively. For *O. nivara*, *O. glumaepatula*, and *O. glabemrrima*, their genome assemblies and annotations were attained from Gramene (http://archive.gramene.org).

### Identification of gene families and phylogenetic tree construction

OrthoFinder version 3.0 was used with default values to sort all protein-coding genes into gene families (Emms and Kelly, 2019). The phylogenetic tree was generated by the built-in Species Tree from All Genes (STAG) algorithm in OrthoFinder with its default settings. This algorithm infers species trees by leveraging the most closely related genes within single-copy or multi-copy orthogroups, thereby reducing the inaccuracy of inference due to the limited availability of single-copy orthogroups obtained from the species. The iTOL tool (https://itol.embl.de/) was employed to visualise the phylogenetic tree. The numbers of gene families that were present in perennial species but lost in annuals were calculated in the R environment by using the ‘setdiff’ and ‘intersect’ functions, and Venn diagrams were plotted using the ggVennDiagram package (Gao et al. 2021).

### Gene functional annotation

Gene function prediction was performed using EggNOG-mapper v2 (http://eggnog5.embl.de) according to protein sequence similarity to homologous genes with known functions (Cantalapiedra et al. 2021). The EggNOG database v5.0 was used for protein sequence searching and the parameters were set as the following: -m hmmer -d Liliopsida --itype proteins (Cantalapiedra et al. 2021). GO and KEGG terms were extracted from the EggNOG annotation results. Pfam domains were fetched using the pfam_scan tool (Finn et al. 2006) with default settings and the Pfam database (http://pfam.xfam.org). Motifs were analysed using the Simple Enrichment Analysis tool in MEME suite v5.5.7 (Bailey et al. 2015).

### Comparative co-expression network analysis

To construct a co-expression network for *O. rufipogon*, publicly available raw RNA-seq transcriptome data were downloaded from the National Centre for Biotechnology Information (NCBI) Sequence Read Archive (SRA) database (Stein et al. 2018; Feng et al. 2023). Reads were mapped to the *O. rufipogon* reference genome (Li et al. 2020) and processed to obtain the counts per million (CPM) gene expression values using HISAT2 and Subread (Liao et al. 2013; Kim et al. 2019). The resultant expression matrix with 37,590 genes were used to construct the gene co-expression network based on the graphical Gaussian model approach (Ma et al. 2007). Partial correlation coefficients were calculated using GeneNet between all genes and a threshold of 0.0085 was selected (Schäfer and Strimmer 2005). The MCL algorithm (Enright 2002) was employed to cluster the network into modules, which were visualized using Cytoscape version 3.8 (Shannon et al. 2003). For comparative analysis, an *O. sativa* co-expression network (containing 1286 modules) constructed using the same approach was employed (Zhang et al. 2022b). To ensure legitimate comparative analysis, we applied our pipeline to construct a network with the same raw RNA-seq data used in Zhang et al. (2022b), and we obtained identical network results. Nodes from the *O. sativa* modules were mapped to the *O. rufipogon* network using JCVI with parameters ‘-m jcvi.compara.catalog ortholog --cscore = .99’ (Tang et al. 2024). GO and rice TO annotations were downloaded from the Oryzabase database on March 5, 2025 (Kurata and Yamazaki 2006), and enrichment analyses were performed as described in Zhang et al. (2022a).

### Evolutionary analysis

The amino acid sequences of candidate *O. rugipogon* genes were first searched against the genome assembly of *O. nivara* using tBLASTn with default parameters (Camacho et al. 2009). The alignment results were then imported into PseudoPipe for pseudogene identification (Zhang et al. 2006). The R package ‘circlize’ was employed to visualise the location of the identified pseudogenes and their corresponding functional genes in chromosomes (Gu et al. 2014). For Ka/Ks ratio calculation, the amino acid sequences of the pseudogene and functional gene pair were first aligned and then transformed into CDS alignments using ClustalW2 software (Larkin et al. 2007). Subsequently, the alignment results were fed into KaKs_Calculator 3.0 to calculate nonsynonymous and synonymous substitution rates (Zhang 2022b).

## List of abbreviations

CPM: Counts per million
GO: Gene ontology
IPPT: Isopentenyl pyrophosphate transferase
KEGG: Kyoto Encyclopedia of Genes and Genomes
NCBI: National centre for biotechnology information
SRA: Sequence read archive
STAG: Species tree from all genes

## Acknowledgements

We thank Prof. Shilai Zhang from Yunnan University for providing assistance in computational facility usage.

## Author Contributions

Z.Li, F.H., and L.E.R designed and supervised the study. Z.Lu, Y.Z., G.Y., and Z.Li performed the analyses. Z.Li and G.Y. wrote the manuscript with additional inputs from all the authors. All authors read and approved the paper.

## Data Availability

All data supporting the conclusions of this article are provided within the article or in the supplementary material.

## SUPPLEMENTARY DATA

**Supplementary Table 1.** Gene families obtained in this study.

**Supplementary Table 2.** EggNOG annotation of the identified candidate genes.

**Supplementary Table 3.** Enrichment analysis of the identified candidate genes.

**Supplementary Table 4.** Motif enrichment analysis of the identified candidate genes.

**Supplementary Table 5.** Pfam analysis of the identified candidate genes.

**Supplementary Table 6.** Gene co-expression network of *O. rufipogon*.

**Supplementary Table 7.** Module 1 of the *O. rufipogon* network and its connected *O. sativa* modules.

**Supplementary Table 8.** Module 2 of the *O. rufipogon* network and its connected *O. sativa* modules.

**Supplementary Table 9.** Genomic location of the identified pseudogenes in *O. nivara*.

